# The *C. elegans* gonadal sheath Sh1 cells extend asymmetrically over a differentiating germ cell population in the proliferative zone

**DOI:** 10.1101/2021.11.08.467787

**Authors:** Xin Li, Noor Singh, Camille Miller, India Washington, Bintou Sosseh, Kacy Lynn Gordon

## Abstract

The *C. elegans* adult hermaphrodite germ line is surrounded by a thin tube formed by somatic sheath cells that support germ cells as they mature from the stem-like mitotic state through meiosis, gametogenesis and ovulation. Recently, we discovered that the distal Sh1 sheath cells associate with mitotic germ cells as they exit the niche (Gordon et al., 2020). Here we report that these sheath-associated germ cells differentiate first in animals with temperature-sensitive mutations affecting germ cell state, and stem-like germ cells are maintained distal to the Sh1 boundary. We analyze several markers of the distal sheath, which is best visualized with endogenously tagged membrane proteins, as overexpressed fluorescent proteins fail to localize to distal membrane processes and can cause gonad morphology defects. However, such reagents with highly variable expression can be used to determine the relative positions of the two Sh1 cells, one of which often extends further distal than the other.

## Introduction

The *C. elegans* hermaphrodite gonad is a fruitful system in which to study organogenesis, meiosis, and stem cell niche biology. Recent work from our group (Gordon et al., 2020), used two endogenously tagged alleles of genetically redundant innexin genes *inx-8* and *inx-9* to visualize the somatic gonadal sheath of the *C. elegans* hermaphrodite. We discovered that the distal most pair of sheath cells, called Sh1, lies immediately adjacent to the distal tip cell (DTC), which is the stem cell niche of the germ line stem cells. Previously (based on electron microscopy and on cytoplasmic GFP overexpression from transgenes active in the sheath (*lim-7p::GFP)* (Hall et al., 1999) or its progenitor cells (*lag-2p::GFP*) (Killian and Hubbard, 2005)), Sh1 cells were thought to associate only with germ cells well into the meiotic cell cycle, so our finding required a reimagining of the anatomy of the distal gonad.

Here, we confirm that the Sh1 cells fall at the boundary of a population of germ cells in a stem-like state, report other markers that label the Sh1 cells, and verify that these markers can be used to assess gonad anatomy without unduly impacting the gonad itself. We also discuss reagents that are not suitable markers of Sh1 cells, including an overexpressed, functional cell death receptor that is used to mark Sh1 in a recent study (Tolkin et al., 2021). Finally, we consider best practices for using endogenously tagged proteins for cell and developmental studies.

## Results

### Distal Sh1 associates with the population of germ cells that differentiate first when progression through mitosis is halted or Notch signaling is lost

Our first experiment addresses in a new way the question of what type of germ cells associate with the distal Sh1 cell. The DTC expresses the Notch ligand LAG-2, which is necessary to maintain the germ line stem cell pool (Henderson et al., 1994). Recent work has shown that the active transcription of Notch targets *sygl-1* and *lst-1* (Lee et al., 2019) and the accumulation of their proteins (Shin et al., 2017) is restricted to the distal-most germ cells, describing a population of stem-like germ cells ∼6-8 germ cell diameters (∼25-35 µm) from the distal end of the gonad. Our recent work (Gordon et al., 2020) reported that the position of Sh1 coincides with *sygl-1* promoter’s activation boundary on one side and the accumulation of the meiotic entry protein GLD-1 on the other, consistent with the hypothesis that the distal edge of Sh1 falls at the boundary of that stem-like cell population, ∼30 µm from the distal end of the gonad.

A similarly positioned stem-like germ cell population was found in earlier work that used temperature sensitive alleles to perturb germ cell fate or progress through the cell cycle. The readout was germ cell fate as determined in one of two ways. Anti-phosphohistone H3 staining of proliferative cells and GLD-1 antibody costaining for cells accumulating meiotic factors shows where cells are dividing and beginning to differentiate, respectively. Alternatively, the “transition zone” in germ cell nuclear morphology between “mitotic” and “meiotic” zones can be visualized by the presence of crescent-shaped nuclei of meiotic prophase observed by DAPI staining. We used this latter method of visualizing nuclear morphology.

We repeated these experiments in strains that have tagged innexins to mark the distal sheath Sh1 cells to ask which population(s) of germ cells are associated with Sh1. Here, we describe the original findings and their interpretations, and then our new findings.

First, an *emb-30* temperature-sensitive allele is known to cause germ cell division to arrest at the metaphase-anaphase transition, thus halting the distal-to-proximal movement of germ cells that is driven by the proliferation of more distal cells (Cinquin et al., 2010). In a wild-type gonad, germ cells differentiate (enter and progress through the meiotic cell cycle) as they move from distal to proximal (Figure 1A, left). In *emb-30(ts)* gonads, a shift to the restrictive temperature causes proliferation to halt and germ cells to remain stationary within the gonad (Figure 1A, right). These cells can now differentiate in place—or remain in the undifferentiated state—depending on their exposure to the stemness cue. Germ cells that remain in the niche at the distal end of the gonad do not differentiate after 15 hours at the restrictive temperature, while more proximal germ cells do differentiate. Nuclear morphology differs between these two regions of the germ line.

**Figure 1.**
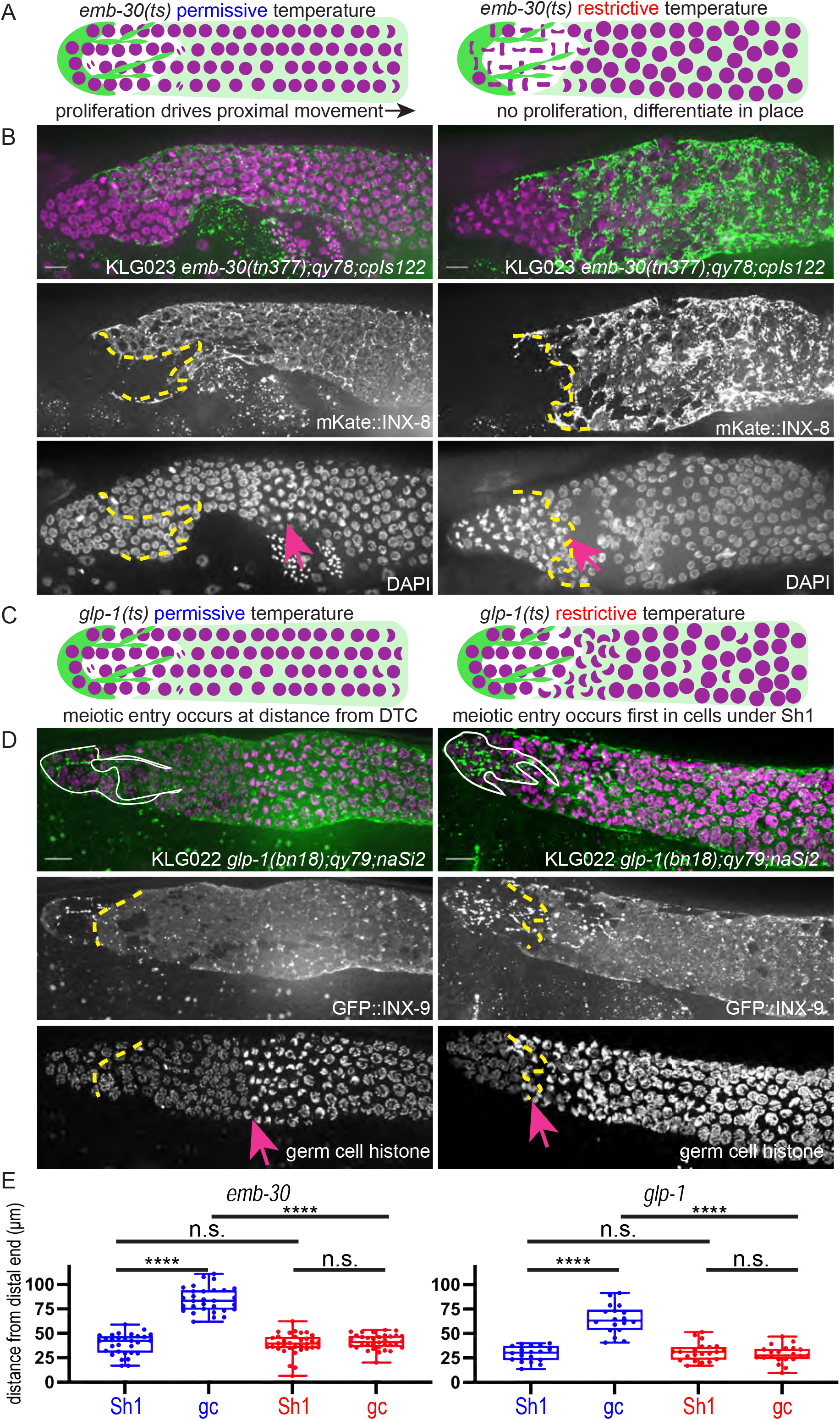
The Sh1 cells associate with proliferative germ cells that are on the path to differentiation. (A) Schematic of hypothesis for *emb-30(tn377)* experiment. Germ cell (gc) nuclei shown in magenta, somatic gonad cells shown in green (DTC) and transparent green (Sh1). (B) Gonads from KLG023 *emb-30(tn377);qy78;cpIs122* worms reared at permissive (left column) and restrictive (right column) temperatures. Top, merged image. Middle, mKate::INX-8 labeling Sh1 (edge outlined with yellow dashed line). Bottom, DAPI staining labeling all nuclei with pink arrow marking gc transition and same yellow dashed line as in middle image showing Sh1 edge. (C) Schematic of hypothesis for *glp-1(ts)* experiment. (D) Gonads from KLG022 *glp-1(bn18);qy79;naSi2* worms reared at permissive (left column) and restrictive (right column) temperatures. Top, merged image. Middle, GFP::INX-9 labeling DTC (outlined in white) and Sh1 (edge outlined with yellow dashed line). Bottom, germ cell histone mCherry (*naSi2[mex-5p::H2B::mCherry]*) with pink arrow showing gc transition and same yellow dashed line as in middle image showing Sh1 edge. Note that the *glp-1(bn18)* allele is not fully wild type at permissive temperatures and is known to have a shortened proliferative zone (Fox and Schedl 2015). (E) Box plots overlaid with all datapoints measuring the distal position of Sh1 and the position of the transition zone in germ cell nuclear morphology. Permissive temperature shown in blue; restrictive temperature shown in red. Permissive *emb-30* N=30; restrictive *emb-30* N=34. Permissive *glp-1* N=18; restrictive *glp-1* N=21. A one-way ANOVA to assess the effect of temperature on proximodistal position of gonad features was performed, and was significant for *emb-30*: F_3,124_=134.5, p<0.0001. Tukey’s multiple comparison test found that the mean values of the positions of Sh1 and the germ cell transition zone were significantly different at the permissive temperature (mean difference of -45.25 μm, 95% CI -52.38 μm to -38.12 μm, p<0.0001), but not at the restrictive temperature (mean difference of -2.30 μm, 95% CI -9.00 μm to 4.40 μm, p=0.808). The position of the germ cell transition zone differed at the permissive vs. restrictive temperatures (mean difference of 42.87 μm, 95% CI 35.95 μm to 49.79 μm, p<0.0001), but the Sh1 position did not (mean difference of -0.078 μm, 95% CI -7.00 μm to 6.84 μm, p>0.9999). This pattern is observed across replicates and various controls (Figure 1—Figure Supplement 1). Similar results were obtained for *glp-1*: F_3,74_=52.84, p<0.0001. Tukey’s multiple comparison test found that the mean values of the positions of Sh1 and the germ cell transition zone were significantly different at the permissive temperature (mean difference of -35.51 µm, 95% CI -44.59 µm to -26.43 µm p<0.0001) but not at the restrictive temperature (mean difference of 2.514 µm, 95% CI -5.892 µm to 10.92 µm, p=0.861). The position of the germ cell transition zone differed at permissive vs. restrictive temperatures (mean difference of 36.02 µm, 95% CI 27.27 µm to 44.77 µm, p<0.0001), but the Sh1 position did not (mean difference of -1.997 µm, 95% CI -10.75 µm to 6.753 µm, p=0.9318). All scale bars 10 µm.

We hypothesized that the transition in nuclear morphology in *emb-30(tn377)* animals would shift proximally after 15 hours at the restrictive temperature (as had previously been observed), to ultimately fall at the distal position of the Sh1 cell as visualized by mKate::INX-8; we hypothesize that the position of the Sh1 cell would itself not be affected by the temperature shift. Indeed, this is what we found (Figure 1B, E), supporting our prior conclusion that there is germ cell fate asymmetry across the DTC-Sh1 boundary. These results are independent of culture time of controls (Figure 1—Figure Supplement 1). The Sh1 cell covers proliferative germ cells outside the niche that are closer to differentiating than those under the DTC.

The second set of experiments using temperature sensitive alleles to reveal differences in germ cell fate along the distal-proximal axis uses *glp-1(ts)* alleles to stop Notch signal transduction and observe where and when germ cells acquire features of differentiation (Figure 1C). The same study (Cinquin et al., 2010) found that the *glp-1(q224)* temperature sensitive allele reared at the restrictive temperature over a 9-hour time course showed a progressively shrinking mitotic zone (as assessed by nuclear morphology of DAPI stained gonads) until hour ∼5.5, at which time the remaining distal most ∼5 rows of germ cells differentiate as a pool. A subsequent study (Fox and Schedl, 2015) used a similar approach (but with the *glp-1(bn18)* temperature sensitive allele, slightly different timing, and antibody staining to determine cell fate) and found a similar result, and additionally discovered that progress through the cell cycle influenced the precise timing of germ cell differentiation.

We used the *glp-1(bn18)* allele to deactivate Notch signaling in a strain with GFP::INX-9 to visualize the Sh1 cells and the fluorescent histone marker *naSi2(mex-5p::H2B::mCherry)* to visualize germ cell nuclei (Figure 1D, E). We hypothesized that after six hours at the restrictive temperature, only the distalmost pool of stem-like cells will not have taken on the crescent-shaped nuclear morphology of meiotic germ cells, while the germ cells under Sh1 will have entered meiosis. We predicted that the Sh1 cell would not change its position across this time interval. Indeed, this is what we found, further supporting our hypothesis that the Sh1-associated germ cells are closer to differentiation than those under the DTC, which are the last to differentiate.

Results from these temperature sensitive mutants confirm what the markers of germ cell fate revealed in Gordon et al. (2020), which is that the Sh1 cell associates with germ cells in the proliferative zone that have left the stem cell niche and are on the path to differentiation, while the stem-like germ cells lie immediately distal to the Sh1 cell at its interface with the DTC.

### Different endogenously tagged membrane proteins reveal a distal position of Sh1

These experiments made use of the endogenously N-terminal tagged *inx-8(qy78[mKate::inx-8])* and *inx-9(qy79[GFP::inx-9])* alleles (Figure 2A and B) generated by (Gordon et al., 2020). Both tagged proteins are highly specific for the somatic gonad throughout development; in the adult, their expression differentiates, with INX-8 signal diminishing from the DTC and INX-9 signal persisting (see white DTC outline in Figure 1D). We have since identified additional endogenous fluorescent-protein-tagged alleles that show expression in the gonadal sheath cells and localize in or near the cell membrane. One of these, *ina-1(qy23[ina-1::mNeonGreen])* (Figure 2C) was briefly reported in (Gordon et al., 2020). We found another that marks the sheath, *cam-1(cp243[cam-1::mNeonGreen])*(Heppert et al., 2018) (Figure 2D). For both tagged innexins as well as *ina-1::mNG* and *cam-1::mNG*, we find that the Sh1 cell has a distal boundary that either displays a measurable interface with the DTC or is so located as to be consistent with such a boundary (where the DTC is not marked by the endogenous protein). The position of this boundary (< 40 µm, or ∼ 8 germ cell diameters) coincides with the domain in which germ cells leave the stem cell niche (Lee et al., 2019) (Figure 2E). We have not yet found a counterexample of an endogenously tagged, membrane-associated protein in Sh1 that demarcates an apparent Sh1 cell boundary at a great distance from the distal end of the gonad in young adults.

**Figure 2.**
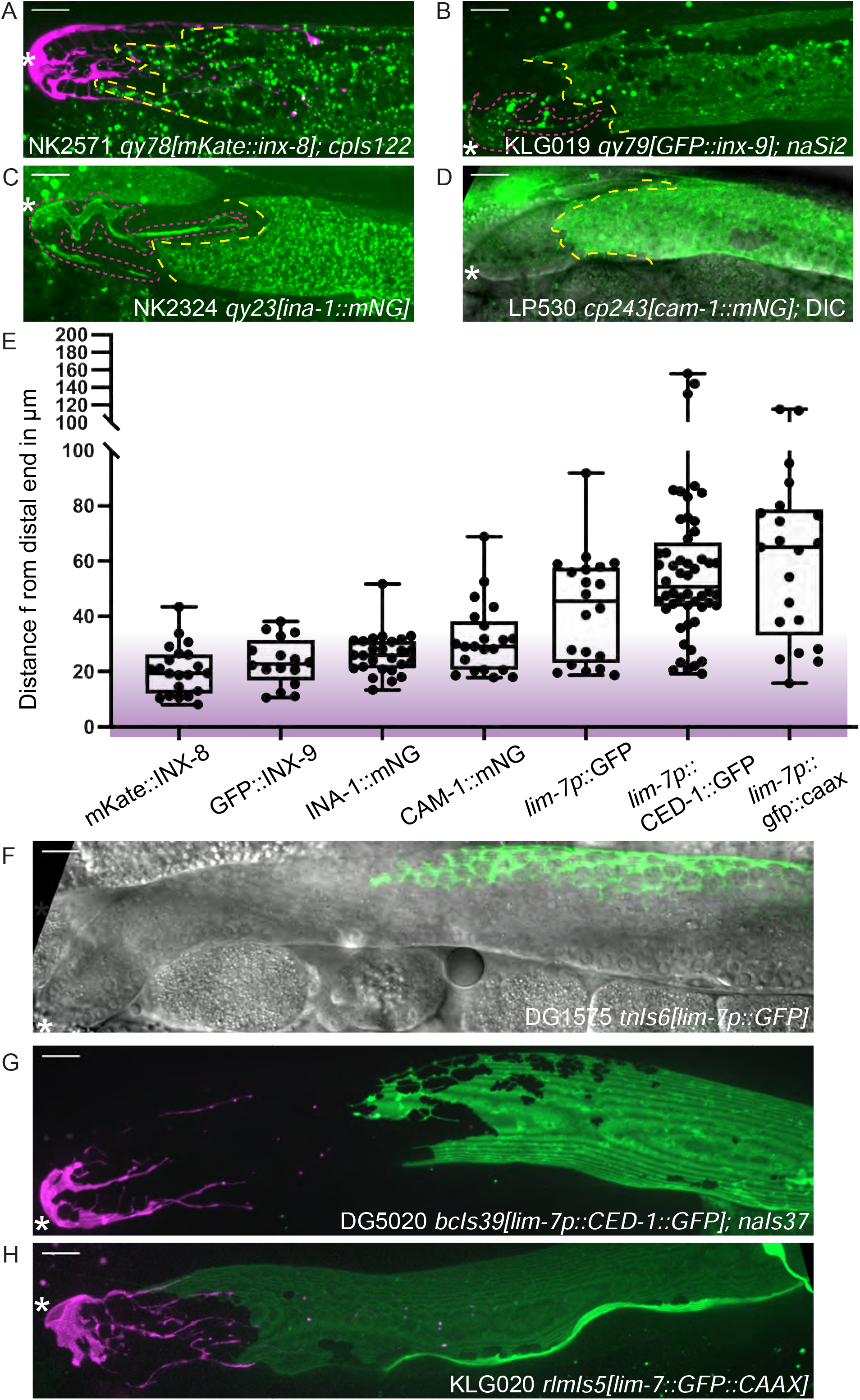
Sheath-expressed fluorescent proteins show consistency among endogenously tagged membrane proteins and greater variability in overexpressed transgenes. (A) NK2571 *qy78[mKate::inx-8]; cpIs122[lag-2p::mNeonGreen:: PLC*^*δPH*^] N=21 (B) KLG019 *qy79[GFP::inx9];naSi2* (channel not shown) N=16 (C) NK 2324 *qy23[ina-1::mNG]* N=26 (D) LP530 *cp243[cam-1::mNG]* N=21 (E) Box plots of Sh1 positions for all strains listed above and below, with fluorescent protein listed on the graph, including transgenes (F) DG1575 *tnIs6[lim-7p::GFP]* N=20 (G) Strain DG5020 *bcIs39[lim-7p::CED-1::GFP]; naIs37[lag-2p::mCherry-PH]* N=52 (note that mean and range agree with those reported in (Tolkin et al., 2021)) (H) KLG020 *rlmIs5[lim-7p::GFP::CAAX]*;*cpIs122* N=21. Purple gradient marks approximate extent of stem cell zone (Lee et al., 2019; Shin et al., 2017).See Figure 2—Figure Supplement 1 for images of minimum and maximum observed distances for all markers. Figure 2—Figure Supplement 2 shows comparisons across development of NK2571 and DG5020. All scale bars 10 µm.

### Overexpressed transgenic markers vary in distal position and expression levels

Three integrated array transgene markers that drive overexpression of fluorescent proteins in the sheath were also analyzed. The first is a *lim-7* promoter-driven cytoplasmic GFP that was used to label the Sh1 cell in a foundational study of the *C. elegans* hermaphrodite gonad, *tnIs6[lim-7::GFP]* (Hall et al., 1999) (Figure 2F). The second is a *lim-7* promoter-driven functional cell death receptor tagged with GFP, *bcIs39[lim-7p::ced-1::GFP]* (Zhou et al., 2001), which is the basis of a recent study that reports a more proximal boundary of Sh1 (Figure 2G, strain DG5020 (Tolkin et al., 2021)). The third is a *lim-7* promoter-driven membrane-localized GFP made by us to mark the sheath cell membrane without tagging an endogenous protein, *rlmIs5[lim-7p::GFP::CAAX]* (Figure 2H). The ranges of the distal edge of GFP localization for all three strains overlaps with what we observed for the four endogenously tagged proteins, but are far more variable, as overexpressed transgenes are known to be (Evans, 2006) (Figure 2E-H, and Figure 2—Figure Supplement 1). Patterns are more similar at earlier developmental stages (Figure 2—Figure Supplement 2).

To untangle this variance, we examined individual worms for evidence of a DTC-Sh1 interface. About half of the scoreable *lim-7p::ced-1::GFP* gonads (strain DG5020) show a DTC-Sh1 interface, and half show a bare region (Figure 3A). We further broke down this dataset by fluorescence intensity of distal CED-1::GFP signal. Strikingly, among animals under a threshold of expression intensity of ∼400 A.U. (less t han 1/3 as bright as the brightest GFP samples), the incidence of a DTC-Sh1 interface was 100% (10/10, as opposed to 15/30 for the whole dataset, Figure 3A). On the other extreme, gonads with stronger CED-1::GFP signal were more likely to have a farther proximal boundary of CED-1::GFP localization. In samples for which CED-1::GFP signal terminates at a great distance from the distal end of the gonad, there are two possible explanations. Either in those animals, the Sh1 position is farther proximal than in animals with other markers, or else CED-1::GFP fails to localize to the edge of the Sh1 cell pair.

**Figure 3.**
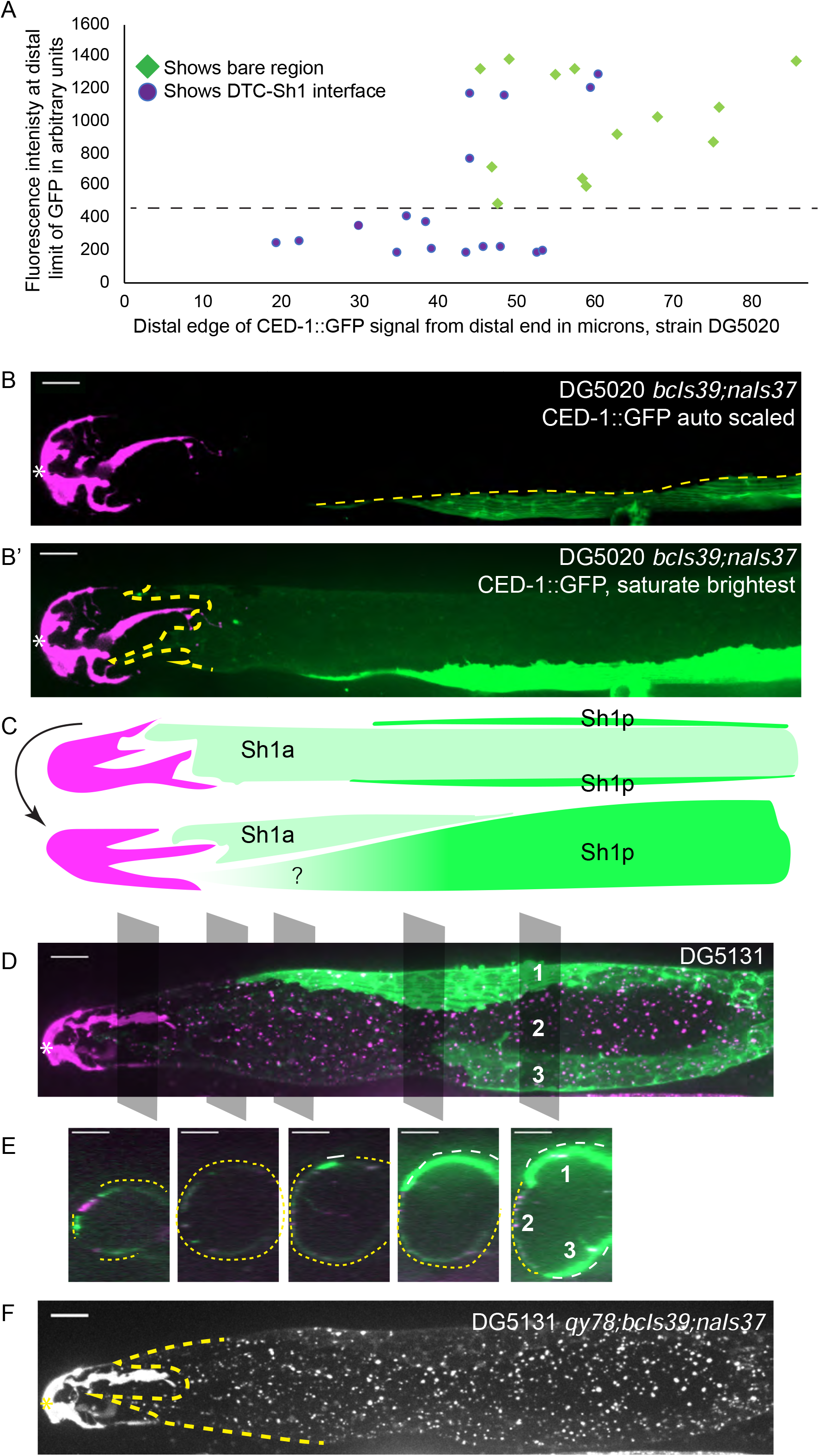
*lim-7p::*CED-1::GFP has variable expression intensity that conceals distal position of Sh1. (A) Plot of distal position vs. fluorescence intensity in arbitrary units of CED-1::GFP at the distal limit of its domain in N=30 DG5020 *bcIs39[lim-7p::CED-1::GFP]; naIs37[lag-2p::mCherry-PH]* animals. Dashed black line: all of the lowly-expressing gonads (under ∼400 A.U., or <30% maximum brightness of brightest sample) have a DTC-Sh1 interface detected. (B) DG5020 sample in which disparate expression levels in the two Sh1 cells of a single gonad arm obscure detection of the DTC-Sh1 interface. The GFP channel is scaled automatically in B; B’ is scaled to saturate the brightest pixels and reveal the dim second Sh1 cell. Dashed yellow link marks the edge of the bright Sh1 cell. (C) Schematic showing Sh1 pair configuration over distal germ line, with the distal extent of Sh1p uncertain in superficial projection. The two Sh1 cells of a pair descend from the anterior and posterior daughters of Z1 and Z4, so the two Sh1 cells are here labeled Sh1a and Sh1p (arbitrarily). Top, superficial view. Bottom, side view. (D) DG5131 *qy78[mKate::inx-8]; bcIs39[lim-7p::CED-1::GFP]; naIs37[lag-2p::mCherry-PH]* sample in which one Sh1 cell contacts the DTC around the circumference of the germ line and the other Sh1 cell lies at some distance from the distal end. Gray boxes and numbers mark planes and landmarks shown in (E). (E) Five cross sections through gonad in (E) made by projecting through two 1 µm re-slices at the positions shown by gray boxes in (D). Same analysis for DG5020 shown in Figure 3 Supplement 1. (F) Same worm as in (D,E); signal from endogenously tagged allele *qy78[mKate::inx-8]* more uniformly labels the Sh1 cells, obscuring their individual shapes. All scale bars 10 µm.

### Expression differences between Sh1 cells in a pair can conceal distal extent of the sheath

We observed a pattern in a subset of gonads where the two Sh1 cells of a pair had dramatically different levels of CED-1::GFP signal, and these cells had different terminal positions on the distal-proximal axis (Figure 3B and 3B’). Exposure time and excitation laser power during image acquisition and subsequent scaling of the resulting image determine whether or not the signal in the lowly expressing cell is readily apparent (Figure 3B vs 3B’). In some cases, the brightness of the other Sh1 cell and the nearby proximal gonad makes the dimmer Sh1 cell nearly impossible to detect. Variable expression levels and even complete silencing of *C. elegans* transgenes are well-known phenomena (Evans, 2006). It was not known, however, that the two Sh1 cells of a pair could assume such different configurations over the distal germline (Figure 3C, and see Figure 2H for the same pattern in the *lim-7p::GFP::CAAX* transgenic strain).

The Sh1 positions become even more clear when *lim-7p::ced-1::GFP* is coexpressed with the mKate-tagged innexin *inx-8(qy78)* in strain DG5131 (Figure 3D and 3E). These markers colocalize in a substantial fraction of animals, as has been reported recently ((Tolkin et al., 2021), see Figure 2 Supplement 2 therein). In the animals that have a discrepancy between GFP and mKate localization in Sh1, the difference in expression reveals an unexpected cell boundary between the two Sh1 cells. We imaged 19 gonads from the coexpressing strain DG5131 through their full thickness. Of those, 4/19 had severe gonad morphology defects (see next section). Of the 15 morphologically normal gonads, 6/15 had discrepancies in CED-1::GFP and mKate::INX-8 signal. In 3/6 such cases, one Sh1 cell makes up the entire DTC-Sh1 interface, with the other terminating at a greater distance from the distal end. In the other 3/6 of cases, both Sh1 cells border the DTC. Fluorescence signal from mKate::INX-8 alone does not allow these cell borders to be detected because that marker is more consistently expressed across the Sh1 cells (Figure 3F).

The variability of the *lim-7p::ced-1::GFP* transgene allowed us to perform something like a mosaic analysis when the two Sh1 cells have very different expression levels but the dimmer cell is still visible (N=31/53 morphologically normal DG5020 gonads imaged to full depth, Figure 3 Supplement 1A-D). Where the borders of the two Sh1 cells can be distinguished, one cell extends at least 20 µm farther distal than the other in 23/31 cases; five additional gonads have expression in only one Sh1 cell that terminates at a great distance (>70 µm) from the distal end. The edges of dimly expressing Sh1 cells can be difficult to resolve. A similar phenomenon was observed when the cytoplasmic GFP of *tnIs6[lim-7p::GFP]* was coexpressed with *qy78[mKate::inx-8]* ((Gordon et al., 2020) Figure 1 Supplement 1 therein). Of note, the N-terminal mKate::INX-8 and GFP::INX-9 tags are most likely extracellular based on the innexin-6 channel structure determined by cryo-EM (Oshima et al., 2016), so there is reason to suspect their localization at the cell membrane will be regulated differently than that of intracellular GFPs.

Additionally, we noticed that in DG5131 gonads where the two Sh1 cells have very different CED-1::GFP expression levels, sometimes mKate::INX-8 is missing from the membrane in Sh1 cells with strong CED-1::GFP signal (Figure 3 Supplement 1E and 1F). Subtracting background, we find that there is a 50% reduction in tagged INX-8 in such membrane regions. Since mKate::INX-8 is a genomically encoded, functional protein, such disruption likely impacts endogenous protein function. This observation hints at a synthetic interaction between the two fluorescent sheath membrane proteins.

### Overexpression of CED-1::GFP transgene is correlated with gonad abnormalities

We therefore asked whether there was further evidence of a synthetic interaction between *lim-7p::ced-1::GFP* and *inx-8(qy78)*. First, we found evidence that suggests that *lim-7p::ced-1::GFP* is damaging to the animals with or without *qy78*. In the strain that expresses *lim-7p::ced-1::GFP* and not *qy78* (strain DG5020), roughly 20% of the animals had profound gonad migration defects in one gonad arm (Figure 4A, 4C). We also observe such defects in the DG5131 strain that combines *qy78[mKate::inx-8]* with the *lim-7p::ced-1::GFP* transgene (Figure 4B, 4/19 or 21% of animals), so we cannot attribute this defect to a spontaneous mutation arising in a single population in transit. We have not observed such morphological defects in the original strain bearing *qy78*, nor in any other strain we have studied. The *lim-7p::ced-1::GFP* transgene seems to cause incompletely penetrant gonad morphology defects.

**Figure 4.**
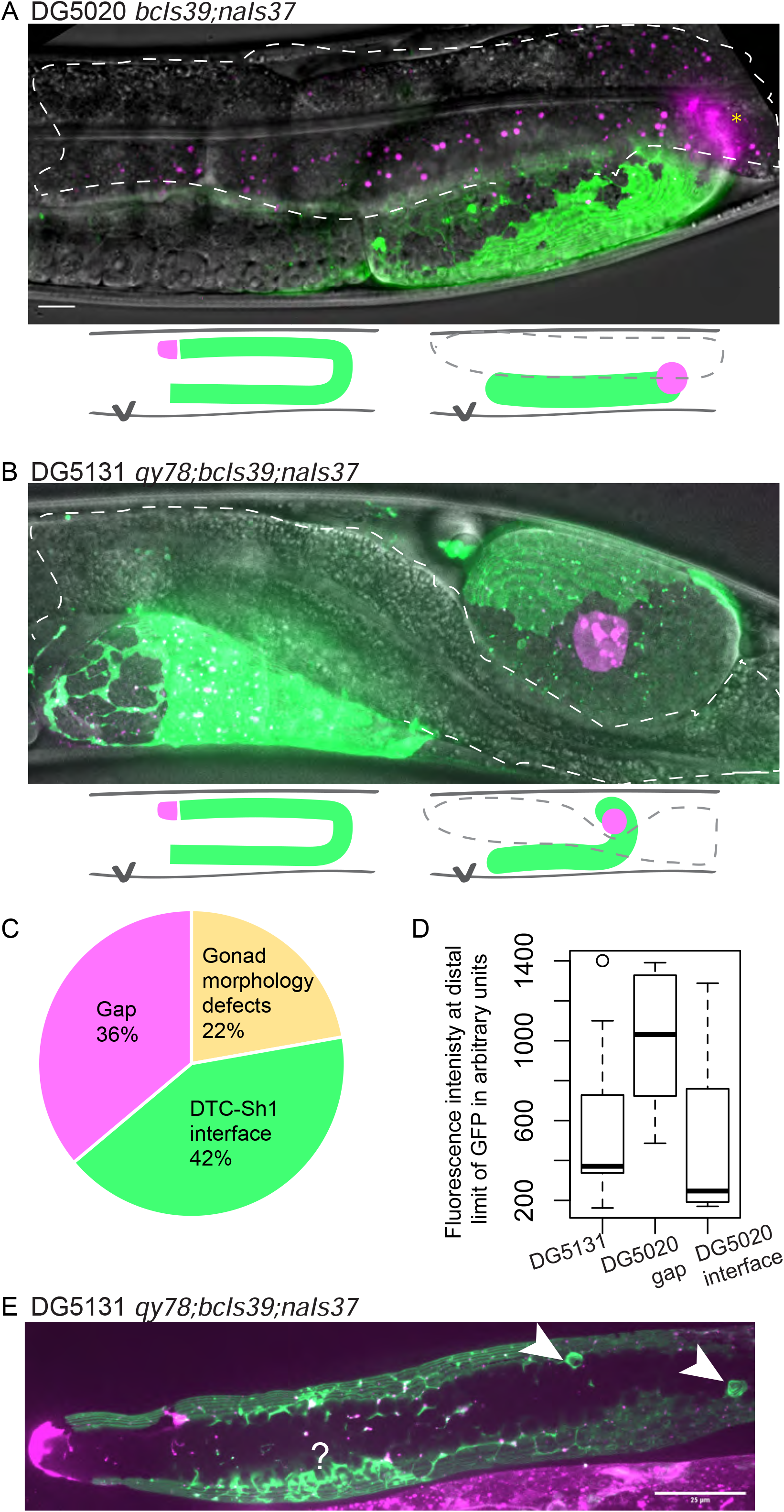
*lim-7p::ced-1::GFP* is correlated with gonad defects. (A) Example of gonad morphology defect in DG5020 *bcIs39[lim-7p::CED-1::GFP]; naIs37[lag-2p::mCherry-PH]* strain, in which the gonad failed to turn. Gut outlined in dashed shape; magenta puncta in that domain are autofluorescent gut granules. (B) Example of gonad morphology defect in DG5131 *qy78[mKate::inx-8]; bcIs39[lim-7p::CED-1::GFP]; naIs37[lag-2p::mCherry-PH]* strain, in which gonad turned once and arrested without elongating along the dorsal body wall. Schematics in A and B show wild-type gonad morphology with two turns and a DTC that arrives at the dorsal midbody, left, beside schematics of defective gonad migration shown in micrographs. (C) Relative proportions of phenotypes observed in DG5020 animals (N=72). (D) Boxplot comparing fluorescence intensity for coexpressing strain DG5131 in addition to data shown in Figure 3 for DG5020. Fluorescence intensity of the *lim-7p::ced-1::GFP* transgene in this background is statistically indistinguishable from expression levels of this transgene in an otherwise wild type background in the subset of samples that display a DTC-Sh1 interface, shown here segregated from samples from this strain that show a gap between the DTC and Sh1 cells. DG5131 N=17. DG5020 gap N=13. DG5020 interface N=17. A one-way ANOVA to determine the effect of category (genotype or presence of an interface) and fluorescence intensity was performed and was significant, F_2,44_=7.70, p=0.001). Tukey’s multiple comparison test finds that the mean fluorescence intensity of DG5020 gonads with a DTC-Sh1 interface differs from DG5020 gonads with a gap between Sh1 and the distal end (p=0.002) and does not differ from DG5131 worms (p=0.908). (E) Gonad from DG5131 strain with white arrowheads indicating aberrant engulfment of germ cells in the distal gonad. Closer to the distal end, a large mass of germ cells showing substantial localization of the CED-1::GFP protein may also reflect ectopic engulfment. Scale bars in A and B, 10 µm; scale bar in E, 25 µm.

Whether or not overexpressed CED-1::GFP also disrupts the localization of untagged innexin proteins or other endogenous sheath membrane proteins as it does the tagged mKate::INX-8, and whether such disruption explains the gonad migration defects we observe for this allele, we currently cannot say. In many of these *qy78; lim-7p::ced-1::GFP* coexpressing animals (strain DG5131), the intensity of CED-1::GFP is notably low (Figure 4D). Lower expression levels of the CED-1::GFP fusion protein, with or without *qy78* in the background, appear more likely to reveal the distal Sh1 cell (Figure 3A and Figure 4D). This could either be because the absence of competing bright signal makes it easier to detect dimly expressing distal Sh1, or because high levels of the transgene product are not tolerated in the distal Sh1 cell. The overexpression of the functional cell death receptor CED-1, and not just the overexpressed membrane-localized GFP, could contribute to the defects observed in this strain. We sometimes observe abnormal sheath membrane protrusions that may result from aberrant engulfment of distal germ cells by the sheath (Figure 4E).

The discrepancy in apparent Sh1 position when two Sh1 cells express different amounts of CED-1::GFP and when CED-1::GFP is coexpressed with mKate::INX-8 provides definitive evidence that CED-1::GFP sometimes fails to label the entire distal sheath (the same phenomenon is reported in Figure 2 Figure Supplement 2B in the recent study (Tolkin et al., 2021)). Furthermore, the defects caused in gonads overexpressing this functional cell death receptor suggests its localization to the distal Sh1 membrane at high levels is not well-tolerated. We therefore conclude that *lim-7p::ced-1::GFP* is an unacceptable marker of distal Sh1.

### Assessing sheath markers for evidence of gonad disruption—Brood size

Just because *lim-7p::ced-1::GFP* is a poor marker of the distal sheath does not, however, relieve concerns that the endogenously tagged innexins mKate::INX-8 and GFP::INX-9 are altering the gonad. A control for tagged innexin function was originally carried out (Gordon et al., 2020). Briefly, a careful genetic analysis (Starich et al., 2014) reported that the single mutant *inx-9(ok1502)* is fertile, but the *inx-8(tn1474); inx-9(ok1502)* double mutant is sterile. Therefore, attempts to use CRISPR/Cas9 to introduce a fluorescent tag in the *inx-8* locus were first performed in the *inx-9(ok1502)* background, and only once a fertile edited strain was recovered was the same edit introduced into the otherwise wild-type genetic background. We conducted brood size assays for strains discussed in this study, including the DG5131 strain containing both *lim-7p::ced-1::GFP* and the tagged innexin *qy78[mKate::inx-8]* that was imaged and analyzed by (Tolkin et al., 2021) but not assayed for brood size (Table 1).

**Table 1.** Brood size assays.

| Strain name | Full genotype | Live brood<sup>a</sup> | Reduction vs wt % | Unhatched eggs | Embryonic Lethality % |
| --- | --- | --- | --- | --- | --- |
| N2 | wild type | 295 ± 39 (n=57) | NA | NA | NA<sup>b</sup> |
| KLG019 | *qy79[GFP::inx-9];nasi2<sup>c\*</sup>* | 226 ± 22 (n=13) | 23% | 165±58 | 41 ± 8% |
| NK2571 | *qy78[mKate::inx-8];cpls122<sup>\*</sup>* | 220 ± 41 (n=15) | 25% | 20±14 | 9 ± 5% |
| DG5020 | *bcls39[lim-7p::ced-1::GFP];nals37<sup>\*</sup>* | 202 ± 29 (n=12) | 32% | 62±47 | 20 ± 14% |
| DG5131 | *qy78[mKate::inx-8];bcls39[lim-7p::ced-1::GFP];nals37<sup>\*</sup>* | 187 ± 45 (n=14) | 37% | 40±25 | 18 ± 11% |
| LP530 | *cp243[cam-1::mNG]* | 260 ± 31 (n=10) | 12% | NA | NA |
| NK2324 | *qy23[ina-1::mNG]* | 237 ± 37 (n=8) | 20% | NA | NA |

<sup>a</sup>Viable offspring that hatch from a single parent
<sup>b</sup>N2 numbers come from multiple trials, not all of which counted negligible numbers of dead embryos, including the trial in which *ina-1(qy23)* and *cam-1(cp243)* were counted.
<sup>c</sup>*qy79[GFP::inx-9]* allele in strains NK2572 and NK2573 from (Gordon et al., 2020) with germ cell nuclear marker *naSi2*; this combination of alleles was used in the cross to *glp-1(bn18)* in Figure 1D.
\*full transgene descriptions in Methods for germ cell (*naSi2*) and DTC (*cpls122*, *nals37*) markers

We find reductions in brood size for all of the strains under investigation, including a reduced brood size and notable embryonic lethality in two strains (DG5020 and DG5131) carrying the *lim-7p::ced-1::GFP* transgene. Interestingly, despite being genetically redundant genes (Starich et al., 2014) tagged in highly similar ways, and having similar live brood sizes, our endogenously tagged *inx-8(qy78) and inx-9(qy79)* strains had dramatically different degrees of embryonic lethality, with *qy79* producing over 150 unhatched eggs per worm. All of the fluorescently marked strains have mildly to moderately reduced brood sizes. On the basis of brood size alone, there is not a strong reason to prefer one of these markers over another.

### Assessing sheath markers for evidence of gonad disruption—Proliferative zone

Because brood size is an emergent property of many gonad, germline, embryonic, and systemic processes (including gonadogenesis, stem cell maintenance, regulation of meiosis, spermatogenesis, oogenesis, metabolism, ovulation, and embryogenesis), defects in brood size are not a direct proxy for dysregulation of the germ line proliferative zone. We therefore turned our attention back to the distal gonad and asked whether the strains with fluorescent sheath markers have abnormalities in several metrics (Figure 5A). The length of the proliferative zones differ among strains (as measured by DAPI staining of germ cell nuclei to detect and measure the length of the germ line distal to crescent shaped nuclei of meiosis I, (Hubbard, 2007)) (Figure 5A, 5C, 5D). The NK2571 strain with the tagged innexin *inx-8(qy78)* and DTC marker has a normally patterned distal germ line (average proliferative zone length of 106 µm, or ∼26 germ cell diameters) that is indistinguishable from wild type N2 (average of 109 µm or ∼27 germ cell diameters Figure 5C, 5C’, and 5D). Excluding worms with gross morphology defects, the DG5020 strain bearing a DTC marker and *lim-7p::ced-1::GFP* has a measurably shorter distal germ line (average of 91 µm, or ∼23 germ cell diameters, Figure 5C” and 5D). In the DG5131 strain that combines these alleles, the distal germ line is notably shortened (average of 79 µm or ∼20 germ cell diameters, Figure 5C”‘ and 5D). This is comparable to the defect caused by the *glp-1(bn18)* allele at the permissive temperature shown in Figure 1E. Abnormal distal gonad patterning provides further evidence that a synthetic interaction between the *lim-7p::ced-1::GFP* transgene and the *qy78* allele—not the *qy78* allele alone—is responsible for the shorter proliferative zone observed for strain DG5131 (Figure 4 from Tolkin et al., 2021).

**Figure 5.**
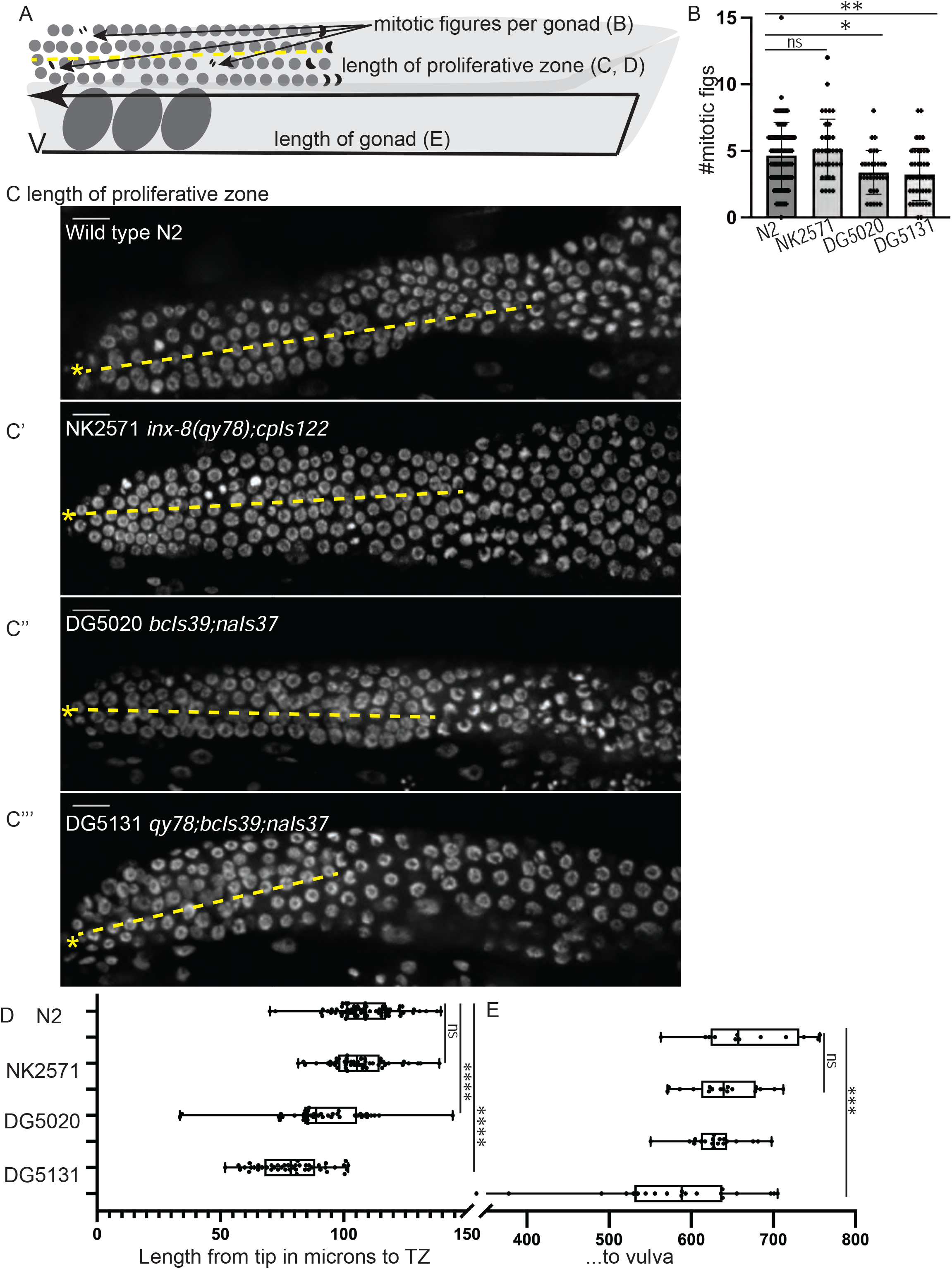
A synthetic interaction between *lim-7p::ced-1::GFP* and the tagged innexin *qy78* shortens the proliferative zone. (A) Illustration of measurements made for Figure 5. (B) Number of mitotic figures observed in DAPI stained animals of the four strains. Numbers of dividing cells and gonads examined are as follows: wild type N2 (N = 311 dividing cells/67 gonads, the NK2571 strain with the tagged innexin *qy78* (N = 184 dividing cells/36 gonads), the DG5020 strain with *lim-7p::ced-1::gfp* (N = 105 dividing cells/31 gonads), the DG5131 strain combining these sheath markers (N= 136 dividing cells/42 gonads). A one-way ANOVA to determine the effect of genotype on number of mitotic figures was significant F_3, 172_ = 7.081, p=0.0002. Tukey’s multiple comparison test revealed that NK2571 did not differ from wild type (mean difference -0.47 cells per gonad, 95% CI -1.65 to 0.71, p=0.7291), DG5020 differed from wild type by 1.26 germ cells per gonad, 95% CI 0.02 to 2.50, p=0.0453), and DG5131 differed from wild type (mean difference of 1.40 cells per gonad, 95% 0.28 to 2.52, p=0.0075). (C-C”“) DAPI stained distal gonads for measurement of proliferative zone for the four strains. Asterisk marks tip of gonad, dashed line marks example of lengths measured. (C) Wild type N2 (N=68), (C’) the NK2571 strain with the tagged innexin *qy78* (N=49), (C”) the DG5020 strain with *lim-7p::ced-1::gfp* (N=40), (C”‘) and the DG5131 strain combining these sheath markers (N=45). Asterisk marks tip of gonad. (D) Plots of proliferative zone length (left) and whole gonad length (right) for the four strains. A one-way ANOVA to determine the effect of genotype on length of proliferative zone was significant F_3,198_=49.15, p<0.0001. Tukey’s multiple comparison test revealed that NK2571 did not differ from wild type (mean difference 2.69 μm, 95% CI -4.283 μm to 9.663 μm, p=0.750), DG5020 differed from wild type by ∼2-5 germ cell diameters (mean difference of 17.94 μm, 95% CI 10.53 μm to 25.36 μm, p<0.0001), and DG5131 dramatically differed from wild type (mean difference of 30.36 μm, 95% CI 23.21 μm to 37.51 μm, p<0.0001). The proliferative zone length of DG5131 was also significantly different from both of its parent strains (NK2571 vs. DG5131 mean difference of 27.67 μm, 95% CI 19.99 μm to 35.35 μm, p<0.0001; DG5020 vs. DG5131 mean difference of 12.42 μm, 95% CI 4.331 μm to 20.50 μm, p=0.0006). (E) Plots of length of entire gonad from tip to vulva. N2 (N=12), NK2571 (N=17), DG5020 (N=19), DG5131 (N=20). A one-way ANOVA to determine the effect of genotype on gonad length was significant F_3,64_=6.27, p=0.0009. Tukey’s multiple comparison test revealed that NK2571 did not differ from wild type (mean difference 31.16 μm, 95% CI -32.99 μm to 95.32 μm, p=0.578), DG5020 also did not differ from wild type (mean difference of 39.42 μm, 95% CI -23.32 μm to 102.2 μm, p=0.3546), and DG5131 did differ from wild type (mean difference of 95.15 μm, 95% CI 33.02 μm to 157.3 μm, p=0.0008). All scale bars 10 μm.

We also counted the number of mitotic figures made by dividing cells in metaphase and anaphase in these strains (Figure 5B) and the total length of the gonad from vulva to tip (Figure 5E). Wild type N2 had an average of 4.6 dividing cells per gonad; NK2571 had an average of 5.1 dividing cells per gonad (these two were not significantly different); DG5020 had an average of 3.4 dividing cells per gonad; DG5131 had an average of 3.2 dividing cells per gonad (these last two strains were significantly different from wild type, see Figure 5B and legend). Gonad lengths were not significantly different between N2 (average length of 670 µm), NK2571 (639 µm) or DG5020 (631 µm), but were significantly shorter in DG5131 (575 µm).

In the end, we find that only the strain combining *inx-8(qy78)* and *lim-7p::ced-1::GFP* has a dramatically smaller gonad that differs from the wild-type in three key measures. Expression of the *qy78* allele alone with a DTC marker, on the other hand, does not cause any of these quantitative gonad phenotypes. The moderate brood size defects shown by all strains could be caused by numerous processes outside of stem cell regulation. For example, we find the hypothesis of (Tolkin et al., 2021) based on the findings of (Starich et al., 2020, 2014), that a major role of *inx-8*/*9* is in the proximal gonad regulating the provisioning of oocytes with essential metabolites, to be compelling.

This hypothesis also has support from the large number of unhatched eggs observed for *inx-9(qy79[GFP::inx-9])*. Thus, we conclude with the observation that endogenous, fluorescently tagged sheath membrane proteins consistently mark both of the distal Sh1 cells without measurably impairing distal gonad function and should be the reagents of choice for live-imaging in this cell type. They also consistently report a distal Sh1 position adjacent to the stem cell zone, as we previously found (Gordon et al., 2020).

## Discussion

We discovered that the distal position of Sh1 is much closer to the distal end of the young adult hermaphrodite gonad then than was previously observed, where it forms an interface with the DTC’s proximal projections and overlaps substantially with the proliferative zone of the germline where mitotic cell divisions occur (Gordon et al., 2020). We have now confirmed this finding with functional manipulations of germ cell cycling and cell fate. We observed a distal Sh1 position in other strains with endogenously tagged sheath cell membrane proteins that act in molecular pathways outside of gap junctional coupling, and in a substantial fraction of traditional transgenic animals expressing *lim-7* promoter-driven CED-1::GFP, GFP::CAAX, and cytoplasmic GFP (though these strains have high variability in fluorescence intensity and localization). Therefore, we consider the results presented here to be confirmatory of the foundational finding of (Gordon et al., 2020), which is that almost all mitotic germ cells in the adult hermaphrodite contact the DTC or Sh1, with a noteworthy population in contact with both. Other recent work suggests a role for the sheath cells in promoting adult germ cell proliferation, specifically through modulation of Notch receptor *glp-1* expression (Gopal et al., 2020). We focus especially on young adults in these studies (less than 24 hours post mid-L4, see Methods). An important caveat to the work is that the gonad is dynamic and cell shapes and positions change over time. Indeed, dynamic processes could lead to the surprising difference in position often seen between the two Sh1 cells in a single gonad arm, with one Sh1 cell growing more actively over germ cells as they leave the niche. The high variability of an overexpressed *lim-7p::ced-1::GFP* transgene has allowed for this surprising discovery, though that variability makes it a poor marker of the absolute position of the Sh1 cells, it sometimes causes gonad defects, and it interacts synthetically with *qy78* to cause gonad defects.

In physics, the observer effect states that it is impossible to observe a system without changing it. In biological imaging in *C. elegans*, this means that we can either observe wild-type animals that are dead, dissected and/or fixed and coated or stained, or we can observe genetically modified animals that are alive. Some fine, membranous cellular structures do not survive fixation (Gerdes et al., 2013; Kornberg and Roy, 2014). On the other hand, any genomic modification runs the risk of altering an animal’s physiology.

We feel most confident examining endogenously tagged gene products in Sh1 for several reasons. First, proteins expressed at physiological levels are less likely to directly damage a cell vs. overexpressed fluorescent proteins (Kintaka et al., 2016). Second the ability to cross-reference among strains with different tagged proteins that act in different molecular pathways allows us to use concordant results in reconstructing cell positions; any single marker may or may not localize to the region of interest, but concordant results among independent experiments help construct an accurate picture of the cell. One factor to consider, however, is that not every endogenously expressed protein is likely to localize evenly across all regions of a cell. We would expect in a large cell like Sh1 that interacts with germ cells in many stages of maturation that some cell-surface proteins would be regionalized. Along those lines, it seems possible that the Sh1 cells might have mechanisms to exclude the cell death receptor CED-1 from the cell membrane domain that contacts proliferating germ cells. The *bcIs39* transgene is typically used to study engulfment of apoptotic germ cell corpses at the bend of the gonad and rescues *ced-1* loss of function mutants for apoptotic germ cell corpse engulfment (Zhou et al., 2001). We find this marker to be unreliable in the distal region of the cell, and to cause gonad defects especially but not only when combined with endogenously tagged *inx-8(qy78)*. A recent study (Tolkin et al., 2021) uses this transgene in most of the backgrounds analyzed (sometimes detecting the CED-1::GFP by anti-GFP antibody staining, which appears to amplify the variability of the marker), so we find this problematic reagent to undermine that study’s conclusions.

The need for caution when observing and interpreting endogenously tagged fluorescent proteins is noted. Several steps can and should be taken to increase confidence that a tagged protein is not causing cryptic or unwanted phenotypes. First, multiple edited lines should be recovered and outcrossed, thereby reducing the likelihood that a phenotype is caused by off-target Cas9 cutting creating lesions in any individual edited genome. Second, brood size should be estimated either by timed food depletion (less rigorous) or formal brood size assays (more rigorous). Third, edited lines should be examined for known phenotypes caused by loss of function of the targeted genes. This can be done, in order of least to most rigorous, by consulting the literature, by comparing to RNAi treatments or known mutants, and finally by introducing AID tags and using the degron strategy to deplete the gene product under the lab’s exact experimental conditions of choice (Zhang et al., 2015), however this step will not work for extracellular tags (because extracellular AID tags are not accessible to TIR1 ubiquitin ligase). Finally, any “markers” used should be assessed on their own for the phenotype of interest. Even with these controls in place, synthetic interactions can emerge between “markers” and alleles, including tagged proteins of interest. These interactions can themselves reveal biologically relevant phenomena, but only if they are recognized.

In the end, no transgenic or genome-edited strain is truly wild type, and it should be our expectation that such strains might be somewhat sensitized as a result. Indeed, the synthetic interaction we document between *lim-7p::ced-1::gfp* and *inx-8(qy78)* suggests that the *qy78* is sensitized for gonad defects caused by other genetic elements affecting the gonadal sheath. However, the perfect reagent does not exit. We can only look for congruent results among a set of independent reagents with non-overlapping weaknesses. Finally, we can formulate questions narrowly enough that, despite their shortcomings, our imperfect reagents are adequate to help answer them. In the future, new endogenously tagged alleles that are expressed in the sheath, single-copy, membrane-localized transgenes that do not affect distal gonad patterning, and different imaging modalities like electron microscopy will shed more light on the complex relationship between the gonadal sheath and the germ line. At the present time, however, we consider the existence of an interface between the DTC and Sh1 cells that coincides with the boundary of the distal-most stem-like germ cells to be supported by the preponderance of evidence.

## Acknowledgements

We thank T. Tolkin, A. Mohammed, T. Starich, T. Schedl, J.A. Hubbard, and D. Greenstein for sharing their manuscript and strains DG5020 (combining published alleles *bcIs39* (Zhou et al., 2001), and *naIs37* (Pekar et al., 2017) and DG5131 (combining published alleles *qy78* (Gordon et al., 2020), *bcIs39* (Zhou et al., 2001), and *naIs37* (Pekar et al., 2017)). We thank D. Greenstein and the CGC for the temperature sensitive mutant strains and B. Goldstein and A. Pani for LP530. We thank R. Dowen and P. Breen for anonymized replication experiments. We are grateful for helpful conversations with D. Sherwood and other colleagues.

## Methods

## Appendix 1. Key Resources Table

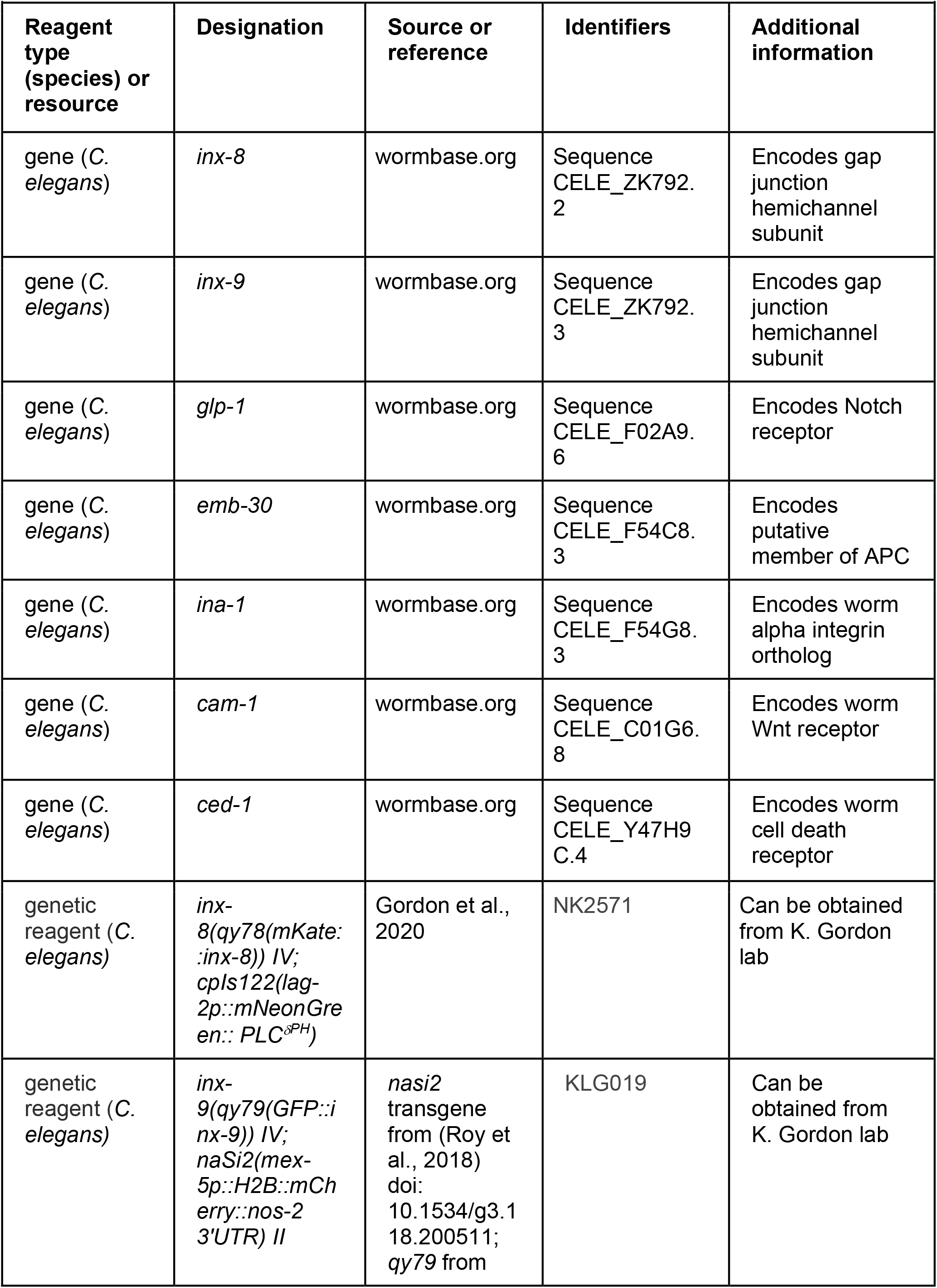
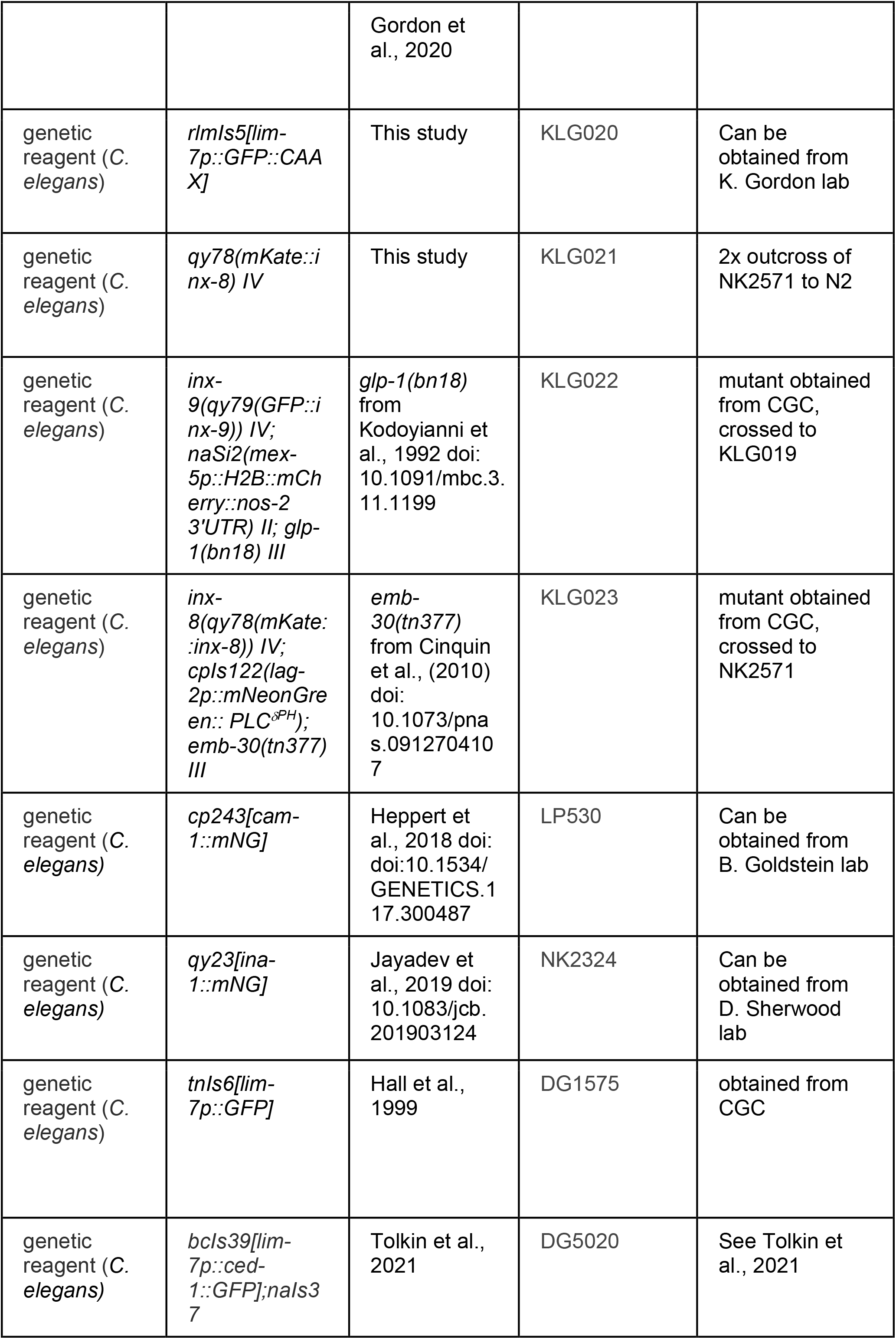
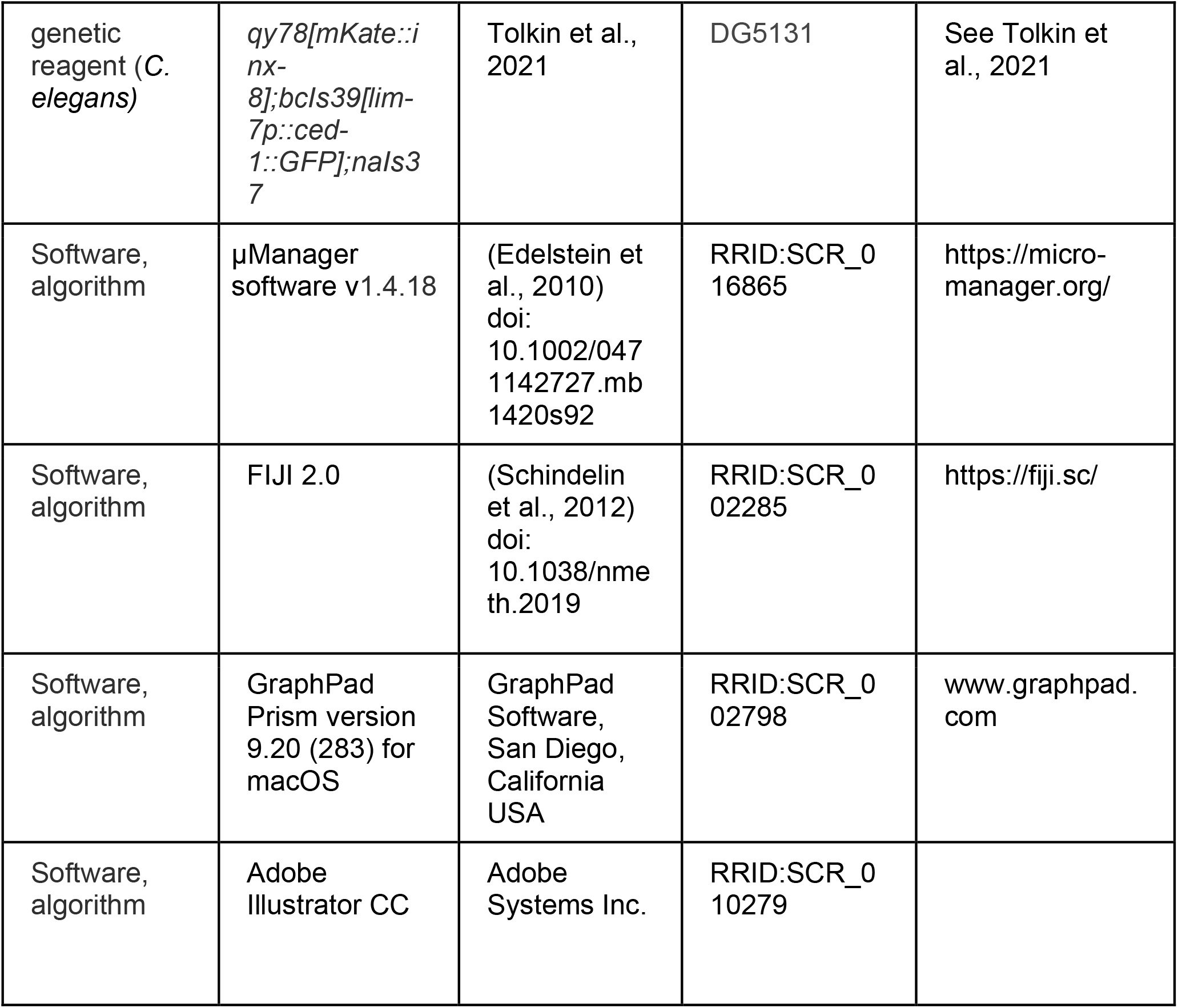

## Strains

In strain descriptions, we designate linkage to a promoter with a *p* following the gene name and designate promoter fusions and in-frame fusions with a double semicolon (::). Some integrated strains (*xxIs* designation) may still contain for example the *unc-119(ed4)* mutation and/or the *unc-119* rescue transgene in their genetic background, but these are not listed in the strain description for the sake of concision, nor are most transgene 3’ UTR sequences.

## Staging of animals for comparisons among sheath markers

We focused on young adult animals around the time egg laying commences, as in (Gordon et al., 2020). Mid L4 animals are picked from healthy, unstarved populations (which are maintained without starving for the duration of the experiment). These animals are kept at 20° C for 16-18 hours, until adulthood is reached and ovulation begins. We prefer not to age the animals much farther into adulthood for routine imaging (though we did this for the temperature shift experiments to follow previously published experimental regimes), as once a full row of embryos is present in the uterus, the distal gonads can become compressed or obscured by embryos. For strains in which a gonad migration defect is observed (DG5020, DG5131), picking animals in the L4 stage prevents bias for or against normal-looking adults (as the defects are profound enough to be visible on the dissecting scope in adults).

## Temperature-sensitive mutant analysis

Worms from the *emb-30(tn377)* mutant genotype were grown at the permissive temperature (16° C) for 24h past L4. Plates were shifted to the restrictive temperature (25° C) for 15 h before DAPI staining, while permissive-temperature controls were maintained at 16° C for 18 h before staining (because development is proportionally slower at 16°C than at 25° C, permissive-temperature controls were cultured longer). Two replicates of this experiment were performed with the results combined in Figure 1E.

Worms from the *glp-1(bn18)* mutant genotype were grown at the permissive temperature of 16° C for 24 h past L4. Plates were shifted to the restrictive temperature (25° C) for 6 h (Fox and Schedl, 2015). Permissive-temperature controls were maintained at 16° C for 6 h. Worms were imaged live (see *Confocal imaging*, below).

## DAPI staining

DAPI staining was by modifying standard protocols (Francis and Nayack, n.d.), with the cold methanol fixation done for a shorter time (2.5 minutes) and the concentration of DAPI higher at 1 ug/ml in 0.01% Tween in PBS in the dark for five minutes, washed once with 0.1% Tween in PBS. Samples were briefly stored at 4° C in 75% glycerol and imaged directly in glycerol solution.

## Confocal imaging

All images were acquired on a Leica DMI8 with an xLIGHT V3 confocal spinning disk head (89 North) with a 63x Plan-Apochromat (1.4 NA) objective and an ORCA-Fusion Gen-III sCMOS camera (Hamamatsu Photonics). RFPs were excited with a 555 nm laser, GFPs were excited with a 488 nm laser, and DAPI was excited with a 405 nm laser. Worms were mounted on agar pads with 0.01M sodium azide (live) or in 75% glycerol (DAPI stained).

## Fluorescence intensity of lim-7p::CED-1::GFP and mKate::INX-8

For quantitative comparisons of fluorescence intensity shown in Figure 3 and Figure 4, gonads were imaged with uniform laser power and exposure times with 1 micron Z-steps. Images were opened in FIJI (Schindelin et al., 2012) and z-projections were made through the depth of the superficial half of the gonad (not including signal from the deep Sh1 cell if it was present). Images without any detectable Sh1 expression were discarded (2/32 images from the analysis in Figure 3A). A line ∼20 µm long parallel to long axis of the gonad, terminating near the distal boundary of GFP expression, and not crossing any gaps in Sh1 revealing background was drawn, and average fluorescence intensity was measured along its length in arbitrary units.

## Measurements of DTC and Sh1 positions

The distal tip of the gonad was identified in the fluorescence images if the DTC was marked or in a DIC image if the DTC was not marked in a given strain. The distance from the gonad tip to the longest DTC process (when marked), and from the gonad tip to the most distal extent of Sh1 was measured in FIJI (Schindelin et al., 2012). A DTC-Sh1 interface is detected by subtracting the first value from the second value—negative numbers reflect the amount of overlap of these cellular domains across the germ line, positive numbers reflect a gap. This is a conservative estimate, as a gap of less than one germ cell diameter (∼5 µm) would still allow germ cells to contact both the DTC and Sh1 at the same time. Min/max settings on the fluorescence images are adjusted to allow the faintest signal to be detected when measuring.

## Analysis of mosaic expression

The variability of the *lim-7p::ced-1::gfp* transgene allowed us to distinguish the two Sh1 cells in a pair, especially when coexpressed with *qy78[mKate::inx-8]*. For this experiment, we imaged animals through the full thickness of the distal gonad (40 µm instead of our usual 20 µm that captures just the superficial half of the gonad that can be imaged more clearly). Animals in which two distinct Sh1 cells had different levels of GFP signal were analyzed further for relative cell position. For DG5131, this was 6/19 samples. For DG5020, this was 31/53 samples.

## Brood size assays

DG5020 and DG5131 were shipped overnight on 9/23, passaged off the starved shipment plate onto fresh NGM+OP50 plates and maintained by passaging unstarved animals for 3 generations before beginning the brood size assay. For each strain, 10-15 L4 animals were singled onto NGM plates seeded with OP50 and kept in the same incubator, on the same shelf, at 20° C. The singled animals were passaged once per day on each of the following 5 days to fresh plates, with all plates maintained at 20° C. Two days after removing the parent, the plates with larval offspring were moved to 4° C for 20 minutes to cause worm motion to cease, and all larvae (and unhatched eggs when noted) were counted on a dissecting scope with a clicker by the same team of worm counters, with internal controls. Plates with unhatched eggs were examined and recounted one day later to see if any hatched. Offspring from parent worms that died or burrowed in the process were not counted. Total sample sizes and results reported in Table 1.

## Distal germ line patterning and mitotic figures

Measurements were made in FIJI from the distal end of the gonad to the transition zone, which is the distal-most row of germ cells with more than one crescent-shaped nucleus. Mitotic figures were counted manually as metaphase or anaphase DAPI bodies. Observations of 0 mitotic figures were counted in the analysis. For Figure 1—Figure Supplement 1E, measurements were made by manually counting cell diameters. In the distalmost region of the restrictive temperature samples, germ cell nuclei are abnormal, so absolute distances in microns were divided by the diameter of a normal-looking germ cell from the distal end to calculate germ cell diameters in this region.

## Gonad length measurements

Strains were synchronized by bleaching and L1 larvae transferred to OP50 seeded NGM plates. At 16C for 48 h. L4 worms were picked to fresh plates and cultured at 16C for an additional 24 hours. These Day 1 adult worms were mounted on agar pads with 0.01M sodium azide and imaged live. Images were analyzed in FIJI using the segmented line tool from vulva to distal gonad tip (usually in 2 tiled images to cover the whole gonad length).

## Statistical analyses

Tests, test statistics, and p values given for each analysis in the accompanying figure legends. One-way ANOVA followed by Tukey’s multiple comparisons test were conducted in R (R Team, 2020) or Prism (GraphPad Prism version 9.20 (283) for macOS, GraphPad Software, San Diego, California USA, www.graphpad.com.

## Figures and Figure Legends

**Figure 1—Figure Supplement 1.**
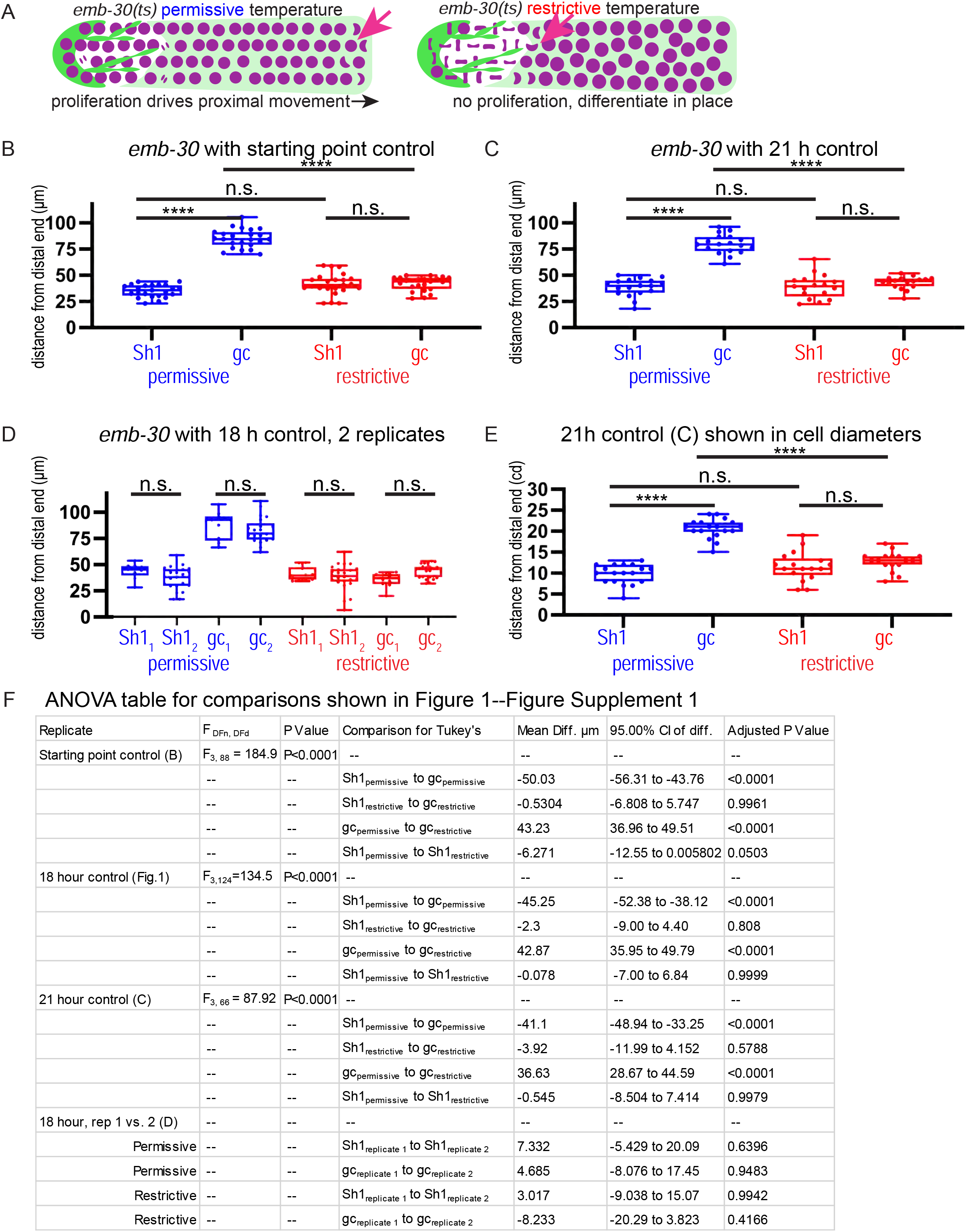
Robustness of *emb-30* temperature shift experimental results to timing of control population. (A) Schematic of hypothesis for *emb-30(tn377)* experiment. Germ cell (gc) nuclei shown in magenta, somatic gonad cells shown in green (DTC) and transparent green (Sh1). (B-F) Results from analysis of strain KLG023 *emb-30(tn377);qy78;cpIs122* under different permissive temperature control culture times. Box plots overlaid with all datapoints measuring the distal position of Sh1 and the position of the transition zone in germ cell nuclear morphology. Permissive temperature shown in blue; restrictive temperature shown in red. (B) Controls fixed at starting point, 36 hours after L4 at permissive temperature as in (Cinquin et al., 2010). Permissive, N=23; restrictive N=23. (C) Controls fixed after 21 additional hours of culture at the permissive temperature. Permissive N=18; restrictive N=17. (D) Two replicates in which controls were cultured an additional 18 hours at the permissive temperature. Replicate 1, permissive N=9; restrictive N=10. Replicate 2, permissive N=21; restrictive N=24. (E) Same experiment as shown in C, but with distances measured in cell diameters instead of microns. (F) Table of relevant ANOVA values for the significance indicators shown in B, C, and D. For (E), a one-way ANOVA to assess the effect of temperature on proximodistal position of gonad features was performed, and was significant F_3,66_=58.44, p<0.0001. Tukey’s multiple comparison test found that the mean values of the positions of Sh1 and the germ cell transition zone were significantly different at the permissive temperature (mean difference of -10.78 cell diameters, 95% CI - 13.09 cd to -8.47 cd, p<0.0001), but not at the restrictive temperature (mean difference of -1.24 cd, 95% CI -3.61 cd to 1.14 cd, p=0.52). The position of the germ cell transition zone differed at the permissive vs. restrictive temperatures (mean difference of 7.73 cd, 95% CI 5.39 cd to 10,08 cd, p<0.0001), but the Sh1 position did not (mean difference of -1.81 cd, 95% CI -4.15 cd to 0.53 cd, p>0.19).

**Figure 2—Figure Supplement 1.**
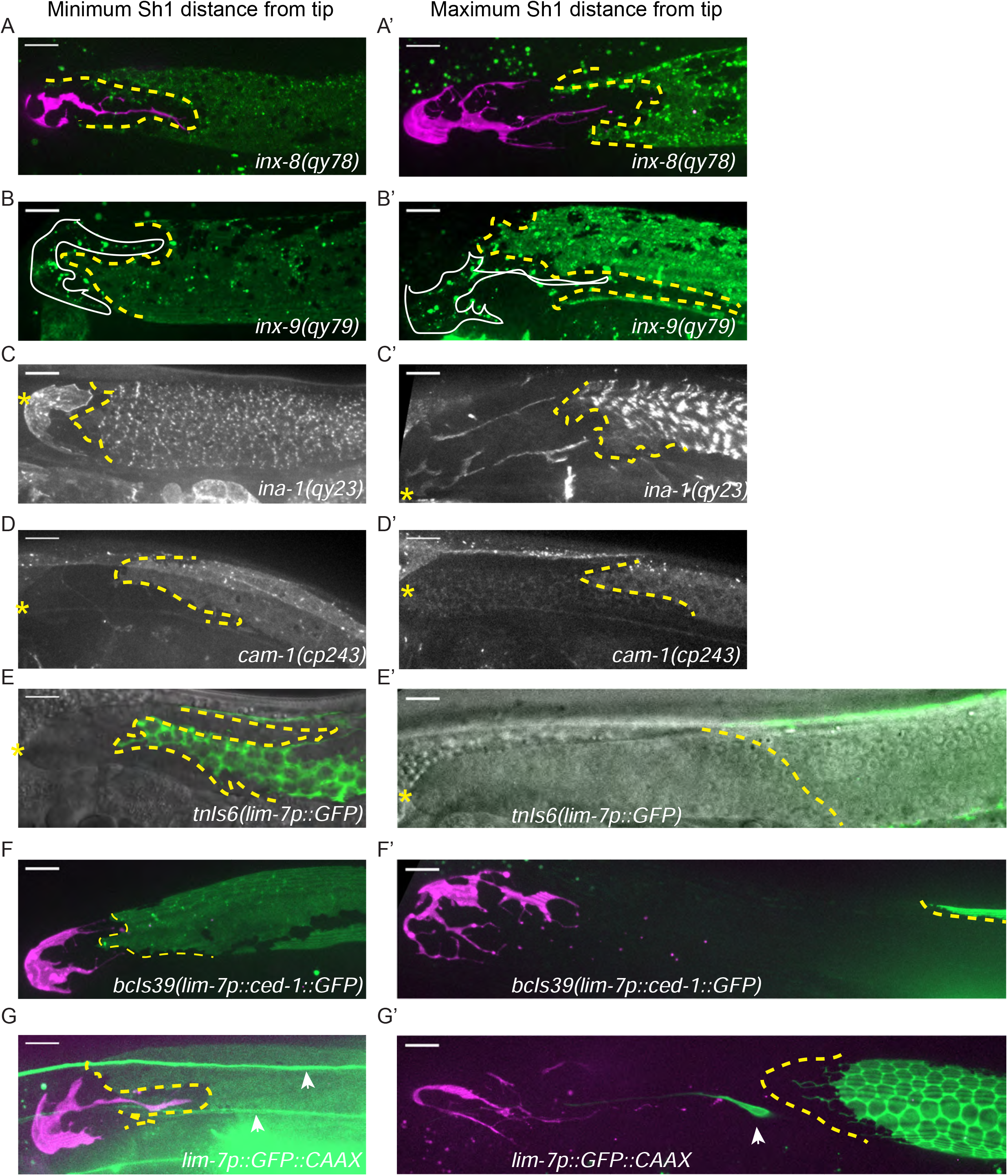
Endogenously tagged fluorescent proteins in the Sh1 membrane are less variable than overexpressed integrated transgenes. Minimum (left column) and maximum (right column) measurements of the distance between distal Sh1 and the distal end of the gonad for (A, A’) *qy78[mKate::inx-8]*, (B, B’) *qy79[GFP::inx9]*, (C, C’) *qy23[ina-1::mNG]*, (D, D’) *cp243[cam-1::mNG]* (E, E’) *tnIs6[lim-7::GFP]*, (F, F’) *bcIs39[lim-7p::ced-1::GFP]*, (G, G’) *rlmIs5[lim-7p::GFP::CAAX]*. Arrowheads in G and G’ mark non-sheath cells positive for *lim-7p*::GFP::CAAX expression in the plane of the gonad and are unavoidably included in Z-projections that capture the gonadal cell surface. Note especially in G how dim the Sh1 expression is at the distal extent, resembling what is sometimes seen for CED-1::GFP expression as reported by Figure 2 Figure Supplement 2B of (Tolkin et al., 2021). In some cases, the selected images are near-minimum or near-maximum due to imaging artifacts like low illumination or sample movement in the true minimum or maximum images. All scale bars 10 µm.

**Figure 2—Figure Supplement 2.**
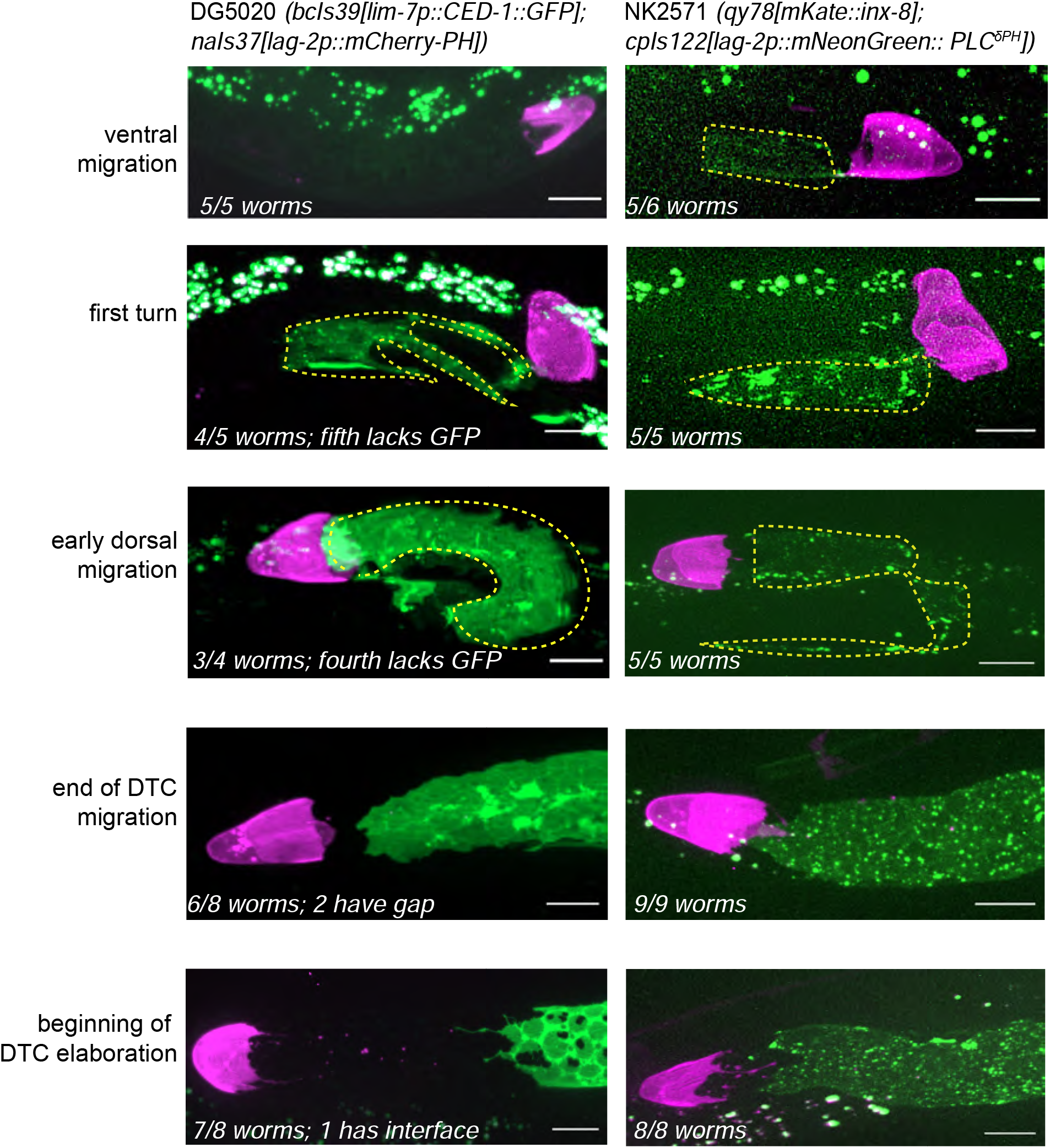
Differences between *qy78(mKate::inx-8)* and *bcIs39(lim-7p::ced-1::GFP)* expression in the sheath appear at the L4-young adult transition. Side by side developmental comparisons between the two favored marker strains. Left column DG5020 (*bcIs39[lim-7p::CED-1::GFP]; naIs37[lag-2p::mCherry-PH]*); right column NK2571 (*qy78[mKate::inx-8]; cpIs122[lag-2p::mNeonGreen:: PLC*^*δPH*^]). Number of worms examined for each strain at each stage given in the figure annotations. Note that both strains show a close association of the DTC and sheath cells during larval development; fluorescence signal from of *lim-7p::ced-1::GFP* appears later in development than *mKate::inx-8* signal (top row). During dorsal DTC migration, highly variable gaps often appear between the DTC and Sh1 in both marker strains (not pictured). Rapid Sh1 growth during this stage has been observed (Gordon et al., 2020), so we attribute these variable gaps to our taking snapshots of a dynamic process. By the end of gonad elongation, most Sh1 cells come to rest within a germ cell diameter of the DTC for both strains. The large gap between the DTC and Sh1 only appears in the *lim-7p::ced-1::GFP* strain after larval gonad migration is complete. This stage is also when the *lim-7p::ced-1::GFP* expressing sheath develops two other unique characteristics—holes over the germ cell bodies, and a frilly distal edge with small, thin distal projections.

**Figure 3 Figure Supplement 1.**
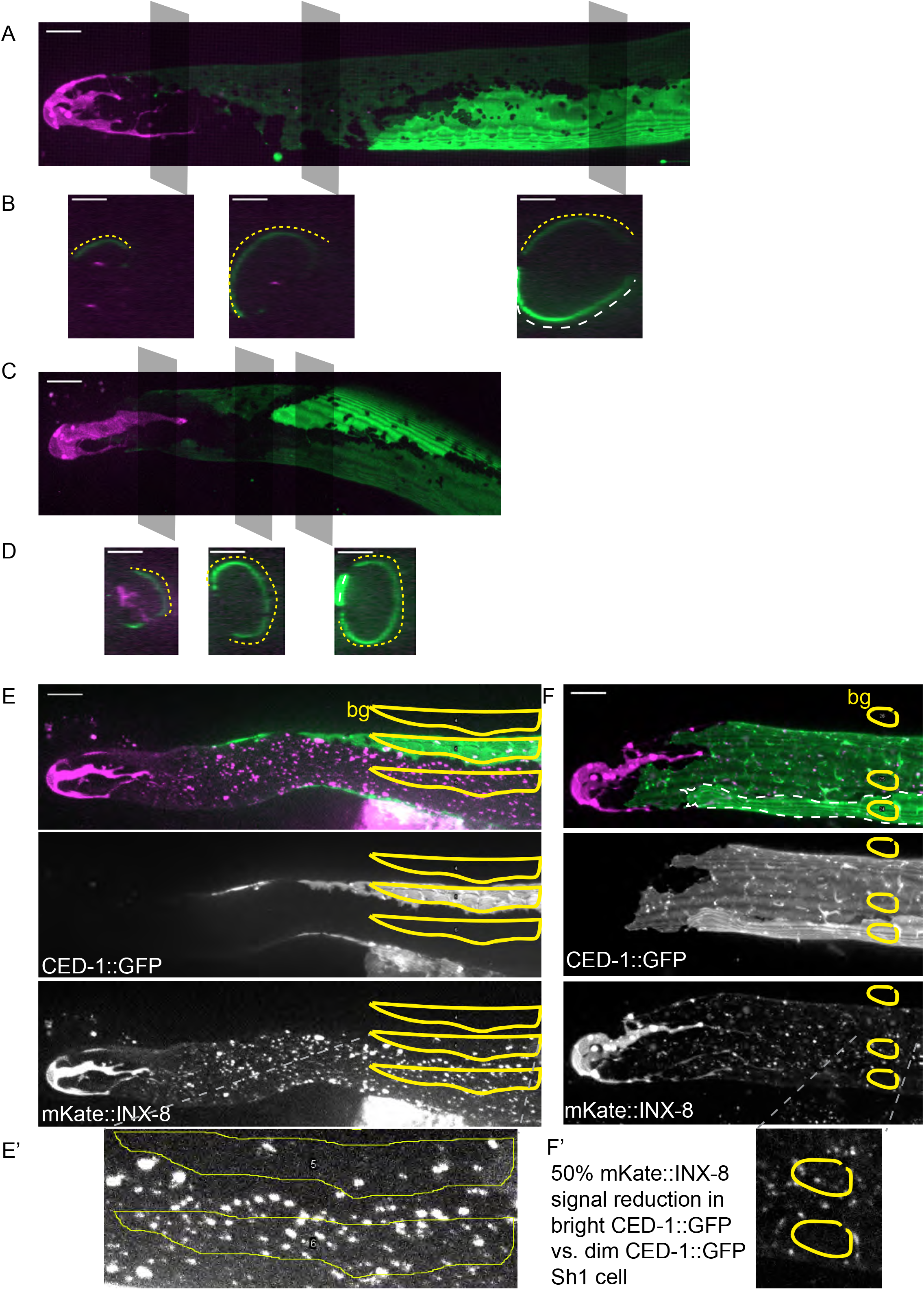
The Sh1 cells of a pair can take two distinct configurations over the distal germ line. (A) Example of a gonad from DG5020 *lim-7p::ced-1::GFP* animal with dramatically different CED-1::GFP signals revealing the shapes of the two Sh1 cells of the pair. Gray boxes show planes depicted in (B). (B) Three cross sections through gonad in (A) made by maximum projection through two 1 µm re-slices (FIJI) at the positions shown by gray boxes in (A). Dashed yellow and white lines mark the two Sh1 cells. Depending on the proximodistal position of the gonad, one or the other Sh1 cell may surround more of the germ line. (C) Example of another gonad from DG5020. (D) Three cross sections through gonad in (C) made by projecting through two 1 µm re-slices at the positions shown by gray boxes in (C). (E,F) Gonads from strain DG5131 with merged images on top, CED-1::GFP channel in the middle, and mKate::INX-8 and distal tip marker channel on the bottom. Yellow outlines show regions of interest in which fluorescence intensity was measured. bg = background, subtracted from fluorescence intensity measured in each of the two Sh1 cells, which express CED-1::GFP at disparate levels. (E’ and F’) Insets from E and F. In both cases, mKate::INX-8 signal is half as strong in the Sh1 cells with more CED-1::GFP. Note also in E and F that mKate::INX-8 and bright CED-1::GFP mark a different distal extent of Sh1. All scale bars 10 µm.

